# Beyond the Plastisphere: Phylogenetic and Biogeographic Diversity of Marine Microorganisms with Putative Plastic-Degradation Potential

**DOI:** 10.64898/2026.09.14.751512

**Authors:** Chloe Kong, Sabrina M. Elkassas

**Author notes:** Address correspondence to Sabrina M. Elkassas, Chloe Kong and Sabrina Elkassas contributed equally to this work.

## Abstract

Plastic pollution is widespread in marine environments, yet the diversity of microorganisms and enzymes that may contribute to plastic degradation remains poorly understood. To identify promising candidates for future experimental studies, we investigated marine microorganisms with proteins related to enzymes previously associated with plastic degradation. Using PlasticDB and the NCBI RefSeq protein database, we identified 523 distinct protein sequences from microorganisms with documented marine provenance. We then examined their evolutionary relationships, environmental origins, and geographic distributions to assess how these candidate proteins are distributed across marine microbial diversity. Only 10.3% of the curated sequence assignments originated from plastisphere or marine-biofilm records, while most were associated with other marine environments, particularly marine hosts. These findings suggest that searches focused exclusively on plastic-associated biofilms may overlook a substantial portion of the marine microbial diversity relevant to plastic transformation. The identified proteins include candidates for direct polyester hydrolysis as well as proteins that may contribute to oxidative chemistry or cellular protection during plastic-associated metabolism. Together, these phylogenetic and biogeographic analyses establish a targeted resource for the discovery and isolation of marine-derived genes with potential roles in plastic transformation, providing experimentally tractable candidates for biochemical characterization and future development of microbial and enzyme-based plastic recycling strategies.

## Importance

Experimental discovery of microbial enzymes for plastic degradation remains relatively low-throughput, despite the increasing urgency of mitigating plastic pollution. Researchers therefore need more targeted ways to identify microbial isolates, genes, and environmental sources that are promising for experimental screening. By using proteins previously associated with plastic degradation to identify related sequences from marine microorganisms, we provide a resource that connects candidate proteins with their evolutionary relationships and environmental origins. This approach helps narrow the experimental search space and guide the selection of candidates for gene isolation and functional testing. We distinguish proteins with plausible roles in direct polymer transformation from those that may instead contribute to oxidative-stress tolerance or persistence in marine biofilms, allowing different protein families to be evaluated using appropriate experimental approaches. Our findings provide a foundation for more targeted discovery of marine-derived enzymes and microorganisms with potential applications in plastic degradation and recycling.

## Introduction

Plastic production has increased rapidly since the 1950s. While ∼21% of the global plastic waste generated is recycled or incinerated, ∼79% still accumulates in landfills or the natural environment; if current production and waste management trends continue, roughly 12,000 Mt of plastic waste are predicted to end up in landfills or the natural environment by 2050 (1). Even if plastic production slows, the most common plastics, such as polyethylene (PE), polypropylene (PP), polyvinyl chloride (PVC), polystyrene (PS), polyethylene terephthalate (PET), and polyurethane (PUR), will persist in the environment because they do not biodegrade; instead, they break down into micro- and nano-plastics that spread through marine, freshwater, and terrestrial ecosystems (1–3).

In marine ecosystems, microplastics affect organisms across vast biological levels, from microorganisms to invertebrates (4). A variety of marine invertebrates readily ingest microplastics, which can be transferred between trophic levels (5, 6). Once ingested, these plastic particles can impede nutrient uptake and cause physical irritation or blockage, compromising the growth and reproduction of these marine organisms (7). Beyond harming marine animals and disrupting entire ecosystems, floating plastic debris at the ocean surface accumulates and converges in each of the five subtropical gyres, forming vast “garbage patches” (8). However, current estimates of the floating plastic at the ocean surface are disproportionately lower than the total wet weight of plastic input over multiple decades; this suggests that a large amount of “missing” plastic is leaving the surface by fragmenting into nanosized particles that are ingested by organisms, then sinking beneath the surface from biofouling or other undetermined reasons (8).

The discrepancy between expected and actual surface plastic loadings demonstrates that current monitoring capabilities are extremely limited, and the plastic pollution problem may be more severe than previously anticipated. In addition to accumulating throughout marine environments, plastic debris creates novel microbial habitats that harbor biofilm-forming communities termed the “Plastisphere” (9). These communities differ from those in the surrounding seawater and have been demonstrated to host microorganisms capable of recycling nutrients and interacting with, transforming, or actively degrading plastic polymers, suggesting that these microbial communities may actively degrade or mineralize synthetic plastic, playing a critical yet poorly understood role in the ocean’s capacity for natural bioremediation (10). Zettler et al. (2013 Zhai et al. (2023) showed that plastic debris hosts a wide variety of microorganisms, including bacteria, fungi, viruses, archaea, algae, and protozoans (9, 11). Among the prokaryotes present were autotrophs, heterotrophs, hydrocarbon-degrading bacteria, and members of *Vibrio*, a genus that includes potential pathogens of humans and marine animals (9, 11). Particularly on floating plastic debris habitats, known as microbial reefs, opportunistic pathogens, such as *Vibrio*, can disperse between ocean basins, potentially introducing harmful microbes into new areas (12). Thus, understanding the biological and physical processes plastics undergo in marine systems and developing solutions to mitigate their ultimate fate in the ocean are imperative.

While examples of solutions degrading plastic waste exist (13, 14), current biotechnological solutions also have limitations (15). Notably, the bacterium *Ideonella sakaiensis strain* 201-F6 can use polyethylene terephthalate (PET), a clear, lightweight thermoplastic polymer resin, as its major carbon source through the combined enzymatic action of polyethylene terephthalate hydrolase (PETase) and mono(2-hydroxyethyl) terephthalic acid hydrolase (MHETase), which degrade PET into terephthalic acid (TPA) and ethylene glycol (13). This finding shows that some microorganisms possess specialized enzymatic pathways capable of depolymerizing synthetic polymers, highlighting the potential of bioremediation. In another recent study by Hachisuka et al. (2023), 91 polyhydroxyalkanoate(PHA)-degrading microorganisms were isolated from enrichment cultures of surface and deep seawater in Suruga Bay (Shizuoka, Japan). 16S rRNA sequencing classified PHA-degrading representatives belonging to *Alloalcanivorax*, *Alteromonas*, *Arenicella*, *Microbacterium*, and *Pseudoalteromonas* (14).

However, both of these observations were made under controlled laboratory conditions, so their relevance to degradation rates in natural marine environments remains uncertain. Additionally, we still lack protein sequences for a majority of these plastic-degrading enzymes and a robust understanding of the global distribution of microbes capable of targeting and degrading the vast majority of plastics produced worldwide. Other than PET and some ester-based polyurethanes, few enzyme systems have been identified that can act on the backbone polymeric components of large-volume-produced plastics. Therefore, while the discoveries of *I. sakaiensis* and PHA-degrading isolates provide strong proof of concept for the microbial degradation of PET and PHA, they represent only a small fraction of the microbial diversity with potential relevance to plastic degradation. Bridging this gap will require not only new enzyme discoveries, but also systematic, data-driven research to identify the diverse range of microbes around the globe with plastic-degrading potential.

To tackle this problem, researchers have created curated databases such as PlasticDB, which exemplify the capabilities of omics studies in exploring microbial plastic degradation (16). As an online platform with an extensive database of microorganisms and proteins involved in the biodegradation of various types of plastic, PlasticDB provides a variety of tools for genome annotation based on published information from peer-reviewed publications (16). Since its launch in 2021, many researchers have begun studying putative plastic-degrading enzymes in the global ocean microbiome (17). A recent 2024 study used PlasticDB’s extensive list of enzymes and other approaches to explore the relationship between the expression of predicted plastic-degradation genes and plastic abundance in the ocean, using data from the Tara Oceans project (18). Database development has advanced significantly in detecting microbial organisms that may carry plastic-degrading abilities, but the evolutionary relationships among putative plastic-degrading microorganisms and their correlation with high plastic-pollution areas remain unclear. By harnessing PlasticDB’s microorganisms and metadata, which contain all current data that have been collated from scientific literature, our study integrates curated genomic databases, comparative sequence analysis, and phylogenetic reconstruction to map the taxonomic diversity and geographic distribution of marine microorganisms with putative plastic-degrading potential.

In this study, we synthesized the published literature on confirmed plastic-degrading enzymes, extracted these enzyme records from PlasticDB, and mined the National Center for Biotechnology Information (NCBI) for homologous gene sequences. From this analysis, we identified candidate plastic-degrading enzymes and their associated microbial taxa. We then reconstructed phylogenetic relationships among these taxa and examined their reported geographic origins to evaluate whether plastic-degrading potential is concentrated within particular evolutionary lineages or marine regions. This framework distinguishes experimentally validated plastic degradation from putative activity inferred from sequence similarity, an important distinction because the presence of an enzyme homolog alone does not demonstrate degradation under environmental conditions. PlasticDB is particularly useful for this purpose because it compiles microorganisms and proteins reported in the literature to be involved in plastic biodegradation, providing a structured starting point for comparative analyses. Ultimately, this study aims to identify taxonomic and geographic patterns that can guide future experimental validation, improving the microbial resources potentially available for plastic-pollution mitigation and providing a catalog of marine plastic-degrading enzymes for future research.

## Materials and Methods

### Dataset Curation and Mining NCBI

PlasticDB (plasticdb.org) summarizes current scientific primary publications on microorganisms and proteomes involved in plastic degradation in a structured format (16). We used PlasticDB’s extensive database, which includes putative plastic-degrading microorganisms collated from scientific literature, to narrow the list down to marine-associated microorganisms with complete protein sequences. We de-duplicated the database and kept only entries associated with marine and marine-associated environments with published protein sequences. The corresponding literature source of each entry was checked for missing data and sequence availability on NCBI. After filtering, we identified nine unique protein sequences with confirmed plastic-degrading potential belonging to the enzymes laccase, medium-chain-length polyhydroxyalkanoate (MCL PHA) depolymerase, oxidoreductase, PETase (2 sequences), PHB depolymerase, glutathione peroxidase (two sequences), and poly(3-hydroxybutyrate) depolymerase (P3HB), as shown in **Table 1**.

**Table 1.** Curated table of solely marine putative plastic-degrading microorganisms and their sequence IDs from PlasticDB.

| Microorganism | Plastic degraded | Enzyme | GenbankID | Reference |
| --- | --- | --- | --- | --- |
| *Psychrobacter* sp. | PE | Laccase | UVG67878.1 | (19) |
| *Pseudomonas alcaligenes* | PHA | MCL-PHA depolymerase | AAQ72538.1 | (20) |
| *Bacillus velezensis* | Polyurethane (PU) | Oxidoreductase | UOM43036.1 | (21) |
| *Marinobacter* sp. | Polybutylene adipate terephthalate (PBAT) and Polylactic acid (PLA) polymer blend (Ecovio-FT) | PETase | UUT36763.1 | (22) |
| *Marinobacter* sp. | PET | PETase | UUT36764.1 | (22) |
| *Alcaligenes faecalis* | PHA | PHB-depolymerase | AAB40611.1 | (23) |
| *Rhodococcus* sp. C-2, | LDPE films | Glutathione peroxidase | WP003945816.1 | (24) |
| *Exiguobacterium marinum* | Additive-free PP film | Glutathione peroxidase | WP026826575.1 | (25) |
| *Shewanella* sp. | P(3HB) | Poly(3-hydroxybutyrate) depolymerase | LC127088.1 | (26) |

Next, to identify additional genes homologous to those with confirmed plastic degradation capabilities, we used BLASTP (27) to query each of the nine candidate plastic-active enzyme amino acid sequences against the NCBI RefSeq protein database (28) using the following parameters: 500 maximum target sequences, E-value of <10^−20^, >70% query coverage, and >40% sequence identity. These parameters provided a systematic screening process across all nine queries such that (**1**) homologs are not included by random chance due to the strict E-value; (**2**) high query coverage prevents partial matches or local domain hits from being mislabeled as full-length functional equivalents; and (**3**) a sequence identity above ∼40% is generally associated with similar 3D structure (29) and function (30) and suggests that putative plastic-degrading genes may also be used for plastic degradation in other organisms possessing the homolog of the gene. Additionally, records described as “partial,” “truncated,” or otherwise incomplete in the BLAST or NCBI protein record were excluded before downstream sequence curation.

Within the top 500 results, every BLASTP hit that met the homology parameters in its original order to determine whether the source organism was isolated in a marine environment. We analyzed each microorganism by inspecting the corresponding accession records (including every linked underlying genome, assembly, strain, and nucleotide record) and available BioProject or BioSample metadata. Since RefSeq WP protein accessions are non-redundant records that may represent identical proteins encoded by multiple strains, assemblies, or genomes, all WP accessions underwent additional review using the NCBI Identical Protein Groups (IPG) report (31). Any sequences whose record description stated they were isolated from freshwater, soil, cropland, rhizosphere, compost, wastewater, sludge, landfill, inland saline, food-associated, or other non-marine environments were excluded from the final curated list. Records with generic, incomplete, ambiguous, unavailable, mixed, or conflicting provenance were classified as uncertain and excluded. We only retained sequences from organisms isolated from a marine environment.

For each BLASTP-confirmed hit, we verified and recorded the NCBI protein accession number, source organism, strain/genome reference paper, isolation environment, and exact isolation location for each curated marine candidate plastic-active enzyme homolog (**Table S1**). Where metadata was incomplete or unavailable, we included the original strain description only when it could be unambiguously linked to the same isolate or genome. We assigned each candidate marine plastic-active enzyme homolog to one of ten environmental groups based on the most specific isolation environment, allowing us to test whether phylogenetically related candidate plastic-active enzyme homologs cluster within particular habitat types. The ten categorical groups were: plastisphere/marine biofilm, marine host-associated, mangrove sediment, polar sea ice/cryosphere, deep-sea/chemosynthetic sediment, intertidal sediment/beach sand, intertidal/coastal seawater, deep seawater, marine sediment, and surface seawater. When an accession could plausibly fit more than one category, the following precedence was used to help maintain consistency across all nine data sets (starting from highest precedence): plastisphere/marine biofilm; marine host-associated; mangrove sediment; polar sea ice/cryosphere; deep-sea/chemosynthetic sediment; intertidal sediment/beach sand; intertidal/coastal seawater; deep seawater; marine sediment; surface seawater. These ten environmental groups allowed us to compare homolog distributions across distinct marine ecological niches, and we used distinct marker shapes for both the global distribution map (**Fig. 1**) and the phylogenetic trees (**Figs. 2-4**) below to maintain consistency and clarity.

**Figure 1.**
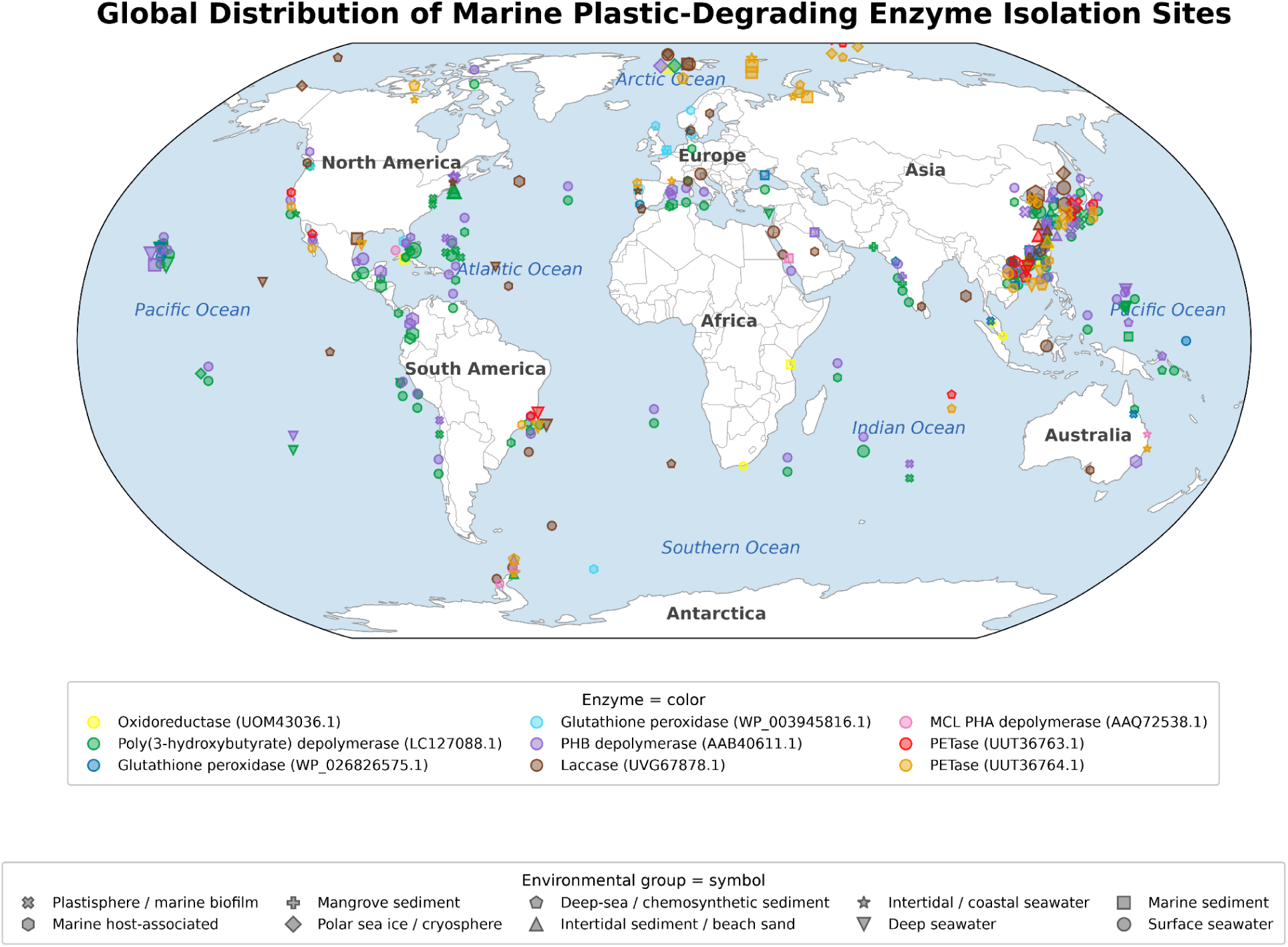
Global distribution of putative plastic-degrading protein homologs recovered from marine-associated microorganisms. Each marker represents the documented isolation location of a microorganism containing a candidate protein homolog included in the nine phylogenetic datasets. Marker color indicates the associated query protein group: oxidoreductase (UOM43036.1), P3HB depolymerase (LC127088.1), glutathione peroxidase (WP026826575.1), glutathione peroxidase (WP003945816.1), PHB depolymerase (AAB40611.1), laccase (UVG67878.1), MCL PHA depolymerase (AAQ72538.1), PETase (UUT36763.1), or PETase (UUT36764.1). Isolates were assigned to standardized marine environmental categories based on available source metadata; locations without reported latitude and longitude were not plotted.

**Figure 2.**
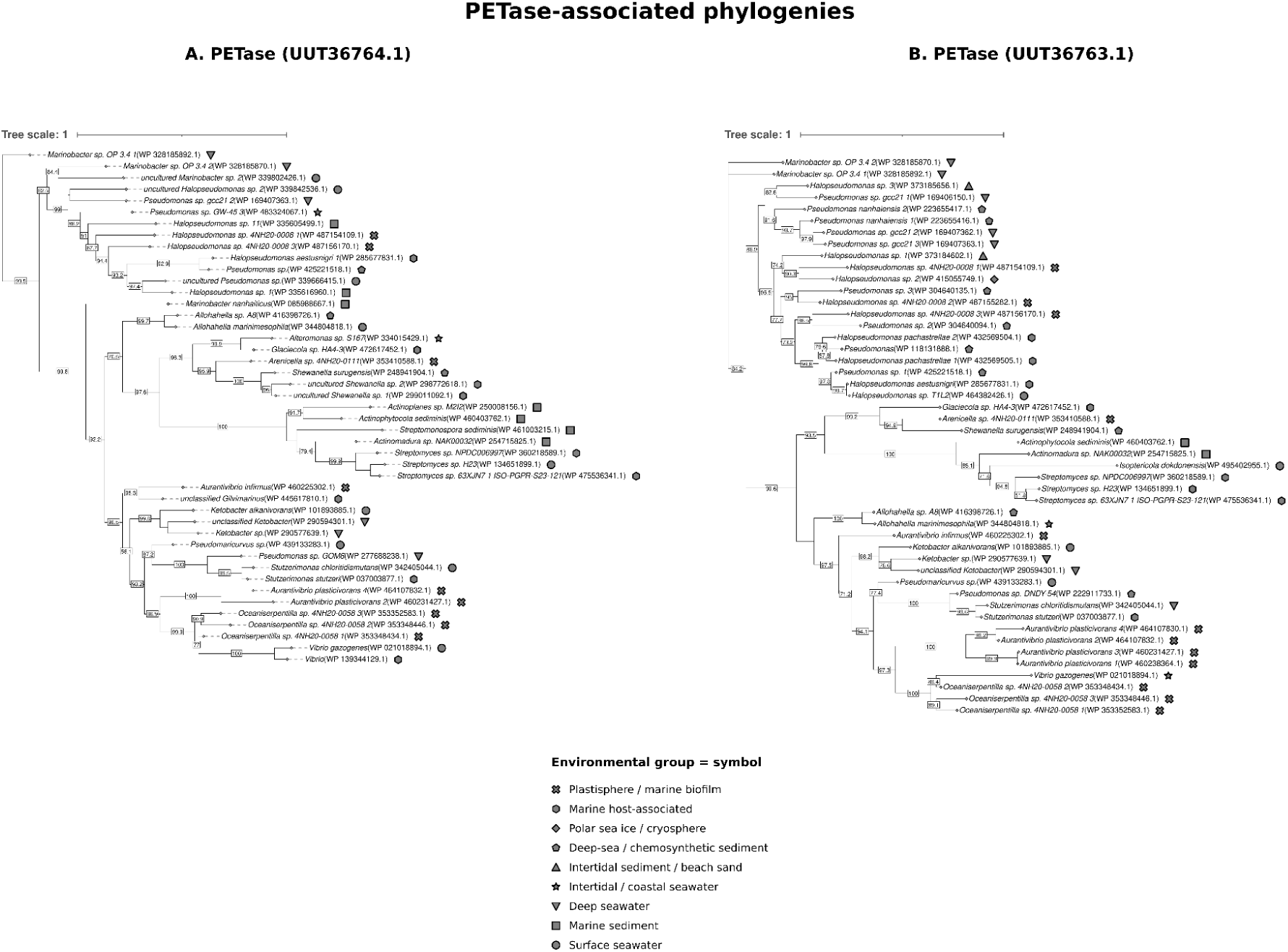
Phylogenetic relationships and environmental provenance of PETase-associated protein homologs. Maximum-likelihood phylogenies of candidate homologs recovered using (A) PETase (UUT36764.1) and (B) PETase (UUT36763.1) as BLASTP query sequences. Terminal labels identify the source organism and NCBI protein accession number; symbols on the right side of the tree tips indicate the verified marine isolation environment associated with each accession. The percentage of replicate trees in which the associated taxa clustered together in the bootstrap test (1000 replicates) is shown at internal nodes; only bootstrap values >50% are displayed. See **Fig. S1** for the complete PETase (UUT36764.1) phylogeny.

We prioritized classifying plastisphere/marine biofilm over the other environmental groups because it directly identifies microorganisms recovered from a plastic-associated substrate, which is particularly relevant to evaluating the environmental distribution of candidate plastic-active enzyme homologs. For example, we classified a plastic-associated sample from mangrove sediment primarily as plastisphere/marine biofilm, while retaining the mangrove setting as supporting contextual information. For every retained accession in the plastisphere/marine biofilm environmental group, we also included the plastic-substrate field when available. We later used the geographic and environmental metadata collected during the BLASTP curation process to visualize the reported distribution of marine candidate plastic-active enzyme homologs.

For each of the nine candidate plastic-active enzyme amino-acid queries, four set of data files were created: a complete FASTA sequence file labeled according to the corresponding confirmed list of marine candidate plastic-active enzyme homologs (see **Supplementary Material**); a metadata table with one row per retained protein accession in the same rank order as the FASTA file (**Table S1**); a geographic metadata file including the protein accession number, latitude, longitude, exact sampling location, coordinate evidence, and the direct source URL **Table S1**); and a file summarizing all key information regarding the entire NCBI BlastP mining process (see **Supplementary Material**).

### Phylogenetic tree

Using each of the nine curated FASTA files containing the marine candidate plastic-active enzyme homolog sequences, we constructed phylogenetic trees. We initially aligned complete homologous amino-acid sequences using MAFFT v7 (32) with the L-INS-i strategy (--localpair --maxiterate 1000). Maximum-likelihood phylogenetic trees were subsequently inferred from amino-acid alignments using IQ-TREE 3 v3.1.3 (33, 34). For each alignment, we selected the best-fitting amino-acid substitution model using ModelFinder (-m MFP) (35). Branch support was evaluated using 1,000 ultrafast bootstrap replicates (-B 1000) (36) and 1,000 SH-like approximate likelihood ratio test (SH-aLRT) replicates (37). For these single-gene trees, clades with UFBoot values of at least 95 percent (36) and SH-aLRT values of at least 80 percent are considered well supported according to the IQ-TREE guidelines (47). We ran IQ-TREE with automatic CPU-thread selection (-T AUTO).

We abbreviated four of the nine phylogenetic trees for figure readability in the main text: The P3HB (LC127088.1), PHB depolymerase (AAB40611.1), laccase (UVG67878.1), and PETase (UUT36764.1) phylogenies contained 188, 167, 140, and 83 protein sequences, respectively. These four larger datasets display a pruned subset of 45 representative taxa rather than every sequence included in the original BLAST-derived dataset. For each reduced tree, we retained all genera represented in the corresponding complete phylogeny, with at least one representative sequence per genus. We allocated additional representatives to more densely sampled genera and selected them to maximize phylogenetic spread across the complete inferred tree, reducing redundancy among closely related sequences while retaining divergent lineages. The full phylogenetic trees are available as panel figures in the **Supplementary Materials** (**Figs. S1-S4**). We visualized and annotated the resulting Newick tree files (.treefile) for all full and abbreviated trees in iTOL v6 (38), with symbols corresponding to the environmental group in the isolation metadata files beside terminal taxa, indicating the verified marine environmental group associated with each sequence. MAFFT and IQ-TREE 3 code and raw .treefiles are available in the GitHub repository: https://github.com/corporeal-snow-albatross-5/Marine_Microbial_Plastic_Degradation_Potential.

### Geographic Distribution Modeling

To assess whether the environmental distribution of the reported plastic-degrading genes in marine microorganisms contained any significant spatial patterns, we developed a custom program, using Python version 3.12.1, integrating geographic sampling record CSV files with standardized environmental metadata Excel files across the nine enzyme datasets: oxidoreductase (UOM43036.1), P3HB depolymerase (LC127088.1), glutathione peroxidase (WP026826575.1 and WP003945816.1), PHB depolymerase (AAB40611.1), laccase (UVG67878.1), MCL PHA depolymerase (AAQ72538.1), PETase (UUT36763.1 and UUT36764.1). We matched geographic and metadata files by NCBI protein accession number and processed them using pandas v3.0.5 (39), NumPy v2.5.2 (40), Matplotlib v3.11.1 (41), Cartopy v0.25.0 (42), and openpyxl v3.1.5 (48) for reading Excel metadata files.

We converted geographic metadata (country, ocean region, or exact GPS coordinates) from the BLASTP analysis to latitude-longitude coordinates, enabling us to plot each isolation location from the list of homologs as color-coded markers for each of the nine candidate plastic-degrading enzyme families. We based each microorganism symbol on the standardized “Environment group” field in the isolation metadata file (**Table S1**), using the same visual encoding scheme. Since different microorganisms were recorded at the same location, we coded custom functions to increase marker size according to the number of repeated records. Marker size increased sub-linearly for repeated observations belonging to the same enzyme, environmental group, and geographic coordinate, in accordance with the equation: *S* = 6 + 2*n* − 1. Since some of the markers were based on multiple sampling coordinate records, we remapped 475 records to 507 points, plotted as location coordinates in the GitHub output file all_mapped_points.csv. The remaining 243 protein accessions that could not be successfully resolved to a plottable location remain as original query rows within all_unmapped_rows.csv. This scaling relationship ensured that repeated markers remained visually apparent while represented sites did not grow too large. Additionally, when overlapping points from different enzyme families were detected, deterministic visual jitter was added to slightly offset the markers from being completely obscured, thereby making the markers in areas of dense sampling more distinguishable.

We constructed the global map using a Robinson projection and simplified it to create a more visually coherent representation. The outputs included one combined map containing all mapped records across the nine query-derived homolog datasets, along with nine individual maps representing each homolog dataset to provide a more detailed view of each enzyme family. However, the geographic data merely describes the reported source-isolate locations and does not represent natural protein abundance, gene prevalence, expression, or experimentally confirmed plastic-degradation activity.

For transparency and reproducibility, all code used in this study to independently model the global distribution of marine-associated plastic-degrading microorganisms is publicly available on GitHub (https://github.com/corporeal-snow-albatross-5/Marine_Microbial_Plastic_Degradation_Potential).

### Environmental distribution analysis

We summarized the environmental distribution of candidate plastic-degradation-associated proteins from the full standardized metadata tables of all nine query-derived homolog datasets. All query-to-hit sequences from **Table S1** were included in this procedure. We first classified each accession into one of the 10 standardized marine environmental categories, then generated a heatmap in R v4.6.1(43) using the tidyverse v2.0.0(44), ggplot2 v4.0.3 (45), and viridis v0.6.5 (46) packages, with each cell showing the within-dataset percentage and raw count (n) for its environmental type. Given that several accessions were shared across multiple query searches, we tallied each accession only once per dataset in which it appeared. Therefore, within each dataset, we calculated the relative abundance of each environmental category as a percentage of the total dataset size rather than the number of distinct proteins discovered. The R script and accession-level metadata, including the exact script and package versions used to generate the environmental-distribution heatmap, are available in the manuscript GitHub repository https://github.com/corporeal-snow-albatross-5/Marine_Microbial_Plastic_Degradation_Potential.

## Results

### Identification and distribution of candidate marine plastic-degradation-associated enzyme homologs

Curation of the PlasticDB Microorganisms and Metadata dataset reduced the initial set of 2,536 records to nine marine-associated protein sequences with complete sequence information (**Table 1**). We used each of the nine protein sequences as a BLASTP query to identify additional homologs from microorganisms with documented marine or marine-associated provenance. These reference proteins represented candidate functions associated with polyester transformation, including PETase-like proteins, PHA-related depolymerases, and oxidative- or stress-associated proteins. The final standardized accession-level metadata contained 718 query-to-hit assignments representing 523 unique RefSeq protein accessions, as some accessions were recovered by more than one query, including two PETase datasets that shared 44 accessions, and the PHB- and P3HB-depolymerase datasets that shared 151 **(Table S1**). Dataset sizes were PETase (UUT36764.1) with 83 accessions; PETase (UUT36763.1) with 48; MCL PHA depolymerase (AAQ72538.1) with 11; PHB depolymerase (AAB40611.1) with 167; P3HB (LC127088.1) with 188; oxidoreductase (UOM43036.1) with 14; laccase (UVG67878.1) with 140; glutathione peroxidase (WP003945816.1) with 22; and glutathione peroxidase (WP026826575.1) with 45. All curated protein accessions were associated with marine bacteria. Ten environmental groups were represented across the nine phylogenetic datasets, which contain genes encoding the enzyme families PETase, laccase, oxidoreductase, glutathione peroxidase, PH3B depolymerase, MCL PHA depolymerase, or PHB depolymerase.

Out of the 718 query-to-hit assignments, 475, corresponding to 337 distinct protein accessions, had data to represent in the global distribution shown in **Fig. 1**. The putative plastic-active enzyme homologs were recovered from microorganisms sampled across the Atlantic, Pacific, Indian, Arctic, and Southern oceans, as well as marginal seas and coastal marine environments, indicating a wide geographic distribution beyond floating plastic debris or surface-water isolates. Along the northwestern Pacific margin comprising the East China Sea, the Yellow Sea, the Korean Peninsula, the coast of Japan, and the South China Sea, lies the most concentrated area of reported isolate records on the global map. These East and Southeast Asia regions appear to contain the greatest concentration of curated records, whereas the sampling distribution across the rest of the globe is more dispersed. These points represent database-linked reported isolation locations, so apparent regional clustering should not be interpreted as true biogeographical hotspots or as an indication of the natural ecological abundance of a specific bacterial lineage.

### Phylogenetic diversity of candidate plastic-active and stress-associated proteins

We created nine phylogenetic trees from the nine candidate plastic-degradation- and oxidative-stress-associated protein homolog datasets in **Table S1**. For PETase (UUT36764.1), PHB depolymerase (AAB40611.1), P3HB depolymerase (LC127088.1), and laccase (UVG67878.1), we displayed a select 45 sequences (see **Materials and Methods**) in the phylogenetic tree panel figures below for readability. The complete phylogenies are shown in **Figs. S1** to **S4**.

Among the two PETase-associated trees, PETase homologs are most abundant among the taxonomic groups of *Marinobacter*, *Halopseudomonas*, and *Pseudomonas*-related spp. (**Fig.2**). The two queries recovered many overlapping protein sequences: both shared 44 accessions between the two datasets, with PETase (UUT36763.1) recovering 83 accessions (**Fig. 2A)** while PETase (UUT36764.1) recovered 48 (**Fig. 2B**). In the PETase (UUT36763.1) tree, *Pseudomonas* and *Halopseudomonas* were the most represented genera, with 10 accessions each, along with several other Actinomycetota genera, including *Actinomadura*, *Actinophytocola*, *Actinoplanes*, *Streptomyces*, and *Streptomonospora*. This broad taxonomy was seen in the PETase (UUT36763.1) phylogeny at greater sampling depth, where *Halopseudomonas* accounted for 25 of 83 accessions and *Pseudomonas* for 17 of 83, with additional *Marinobacter*, *Aurantivibrio*, *Shewanella*, *Streptomyces*, *Oceaniserpentilla,* and other lineages (**Table S1**). Plastisphere/marine-biofilm environmental group representatives appeared in both phylogenetic trees; however, they were among microorganisms isolated from other habitats rather than being restricted to a single plastisphere-specific bacterial lineage.

The PHA, PHB, and P3HB depolymerase-associated tree representatives differed considerably in accession count (**Fig. 3**). MCL-PHA and PHB depolymerase clades contain enzymes that hydrolyze polyhydroxyalkanoates (PHAs; 49, 50); however, the two groups remain phylogenetically distinct. While the MCL-PHA depolymerase dataset formed the smallest subtree with 11 candidate plastic-active enzyme homologs belonging to seven genera: *Pseudomonas*, *Halopseudomonas*, *Zooshikella*, *Atopomonas*, *Parahaliea*, *Pseudomaricurvu*s and *Bdellovibrio* (**Fig. 3A**), both PHB depolymerase and P3HB depolymerase trees displayed at least 45 representatives (**Figs. 3B** and **3C**), with the full trees (**Figs. S2** and **S3**) containing 167 and 188 accessions, respectively, 151 of which were shared. The genus compositions of PHB and P3HB depolymerases were also similar when comparing the relative representation of their most abundant genera, as both had a significant number of representatives from *Pseudoalteromonas*, *Alloalcanivorax*, *Alteromonas*, *Microbulbifer*, and *Rheinheimera*. Additionally, both candidate plastic-degrading enzyme families were represented across all 10 environment categories (**Figs. S2** and **S3**).

**Figure 3.**
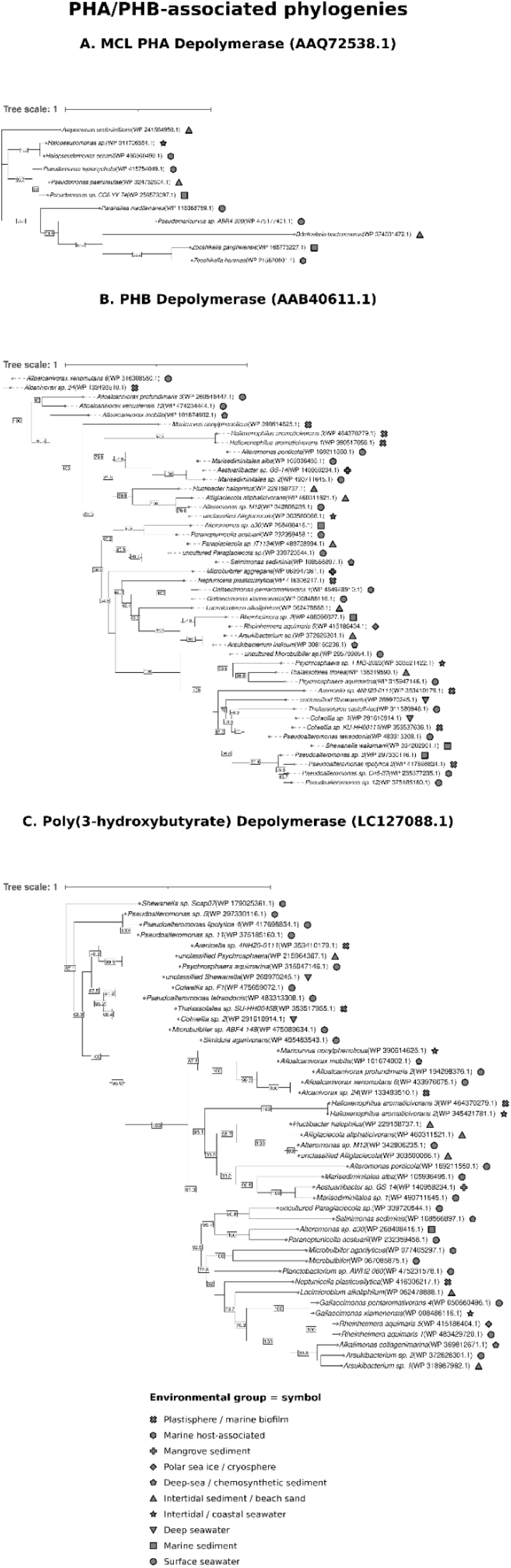
Phylogenetic relationships and environmental provenance of PHA/PHB-associated protein homologs. Maximum-likelihood phylogenies of candidate (**A**) MCL PHA depolymerase (AAQ72538.1), (**B**) PHB depolymerase (AAB40611.1), and (**C**) P3HB depolymerase (LC127088.1) homologs recovered from marine-associated microorganisms. Terminal labels identify the source organism and NCBI protein accession number, and terminal symbols indicate verified marine isolation environments. The percentage of replicate trees in which the associated taxa clustered together in the bootstrap test (1000 replicates) is shown at internal nodes; only bootstrap values >50% are displayed. See **Figs. S2** and **S3** for the complete PHB depolymerase (AAB40611.1) and P3HB depolymerase (LC127088.1) phylogenies.

**Figure 4.**
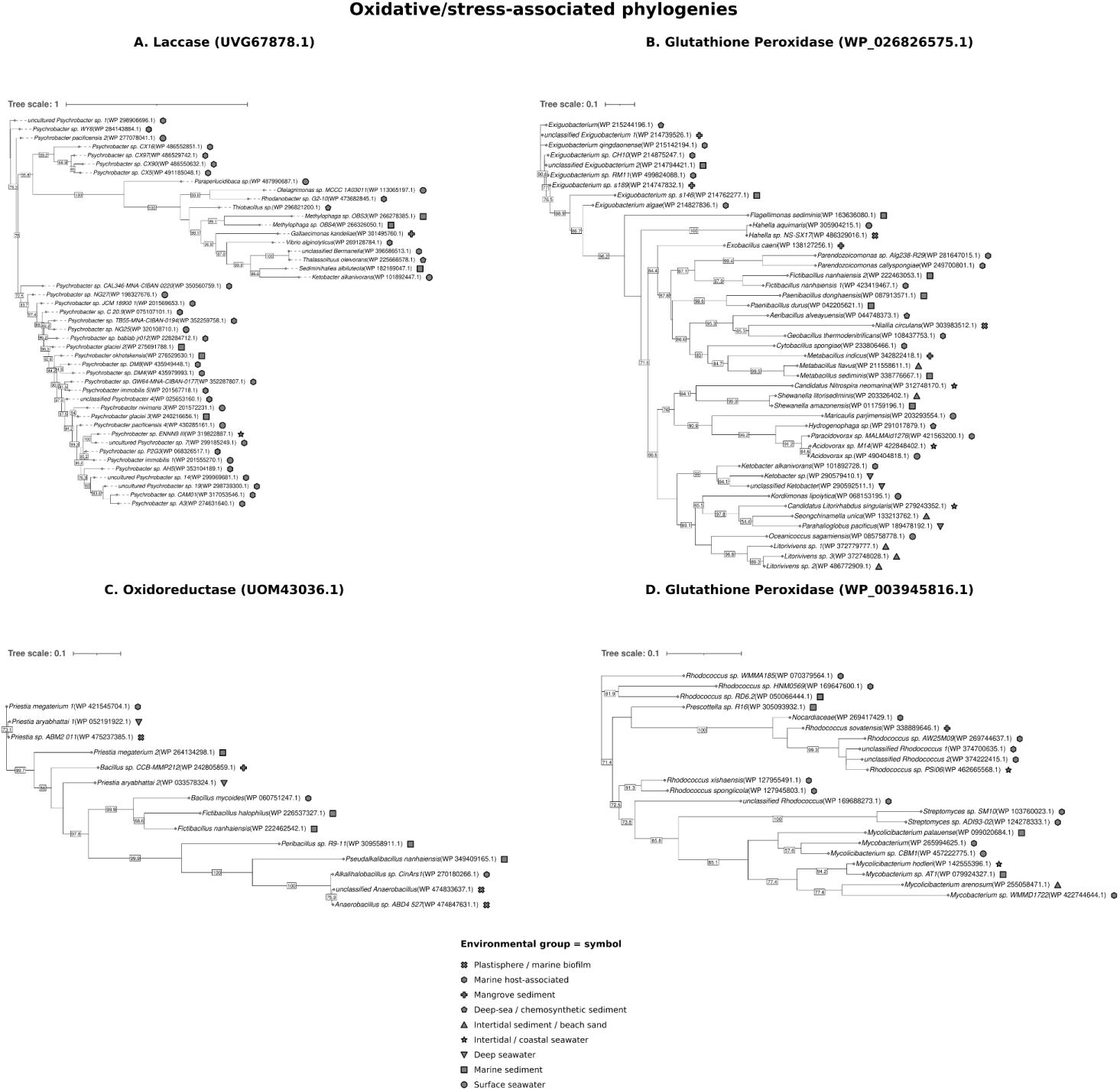
Phylogenetic relationships and environmental provenance of oxidative- and stress-associated protein homologs. Maximum-likelihood phylogenies of candidate (**A**) laccase (UVG67878.1), (**B**) glutathione peroxidase (WP026826575.1), (**C**) oxidoreductase (UOM43036.1) and (**D**) glutathione peroxidase (WP003945816.1) homologs recovered from marine-associated microorganisms. Terminal labels identify the source organism and NCBI protein accession number, while symbols denote verified marine isolation environments. Numbers at internal nodes indicate branch-support values. The percentage of replicate trees in which the associated taxa clustered together in the bootstrap test (1000 replicates) is shown at internal nodes; only bootstrap values >50% are displayed. See **Fig. S4** for the complete laccase (UVG67878.1) phylogeny.

The remaining four trees of the oxidative- and stress-associated datasets showed different taxonomic compositions (**Fig. 4; Table S1**). The dataset encoding for laccase (UVG67878.1) found 140 records, of which 128 were derived from *Psychrobacter* (**Fig. 4A)**. Glutathione peroxidase (WP026826575.1) had 45 accessions (**Fig. 4B)**. The oxidoreductase (UOM43036.1; **Fig. 4C**) and glutathione peroxidase (WP003945816.1; **Fig. 4D**) trees contained fewer representatives. All 14 oxidoreductase (UOM43036.1) homologs belong to bacteria within the phylum Firmicutes (Bacillota), and the associated strains were isolated from deep seawater, marine sediment, mangrove sediment, marine host-associated, and plastisphere/marine biofilm environments (**Fig. 4C**). Glutathione peroxidase (WP003945816.1) contained 22 accessions dominated by *Rhodococcus* (11/22), *Mycolicibacterium* (4/22), and *Mycobacterium* (3/22), with additional *Streptomyces*, *Prescottella*, and *Nocardiaceae* records (**Fig. 4D**). At the same time, Marine biofilm/plastisphere-associated representatives were missing from laccase (UVG67878.1) and glutathione peroxidase (WP003945816.1) trees, but were present in low abundance in the oxidoreductase (UOM43036.1) and glutathione peroxidase (WP026826575.1) trees (**Fig. 4**).

In each of the nine phylogenies, plastisphere/marine-biofilm sequences did not form a single monophyletic assemblage across the putative plastic-active enzyme families; instead, they appeared among phylogenetically related sequences recovered from a variety of non-plastic marine habitats. When grouped by broad functional category, the two PETase-associated phylogenies (**Fig. 2**) accounted for 93 of the 320 displayed representatives (29.1%). The three PHA/PHB-associated depolymerase datasets (**Fig. 3**) accounted for 101 representatives (31.6%), whereas the four oxidative- and stress-associated datasets (**Fig. 4**) accounted for 126 representatives (39.4%). Thus, the candidate dataset included both proteins more directly associated with hydrolysis or transformation of ester-containing polymers and proteins associated with oxidative processes or cellular stress response.

### Habitat-associated phylogenetic patterns from environmental distributions

The environmental composition of all 718 discovered putative plastic-active enzyme homologs in the complete datasets is summarized in **Fig. 5**. Because some accessions appeared in more than one query-derived dataset, the total dataset size used to calculate the percentages was 718 rather than the 523 unique accessions. The largest environmental group was surface seawater with 205 (28.6%), followed by marine host associated isolates (199/718, 27.7%), plastisphere/marine biofilm associated (74/718, 10.3%), marine sediment (56/718, 7.8%), deep seawater (48/718, 6.7%), deep-sea or chemosynthetic sediment (41/718, 5.7%), intertidal sediment or beach sand (35/718, 4.9%), intertidal or coastal seawater (30/718, 4.2%), mangrove sediment (18/718, 2.5%), and polar sea ice or cryosphere (12/718, 1.7%). The values describe only the curated, query-derived datasets.

**Figure 5.**
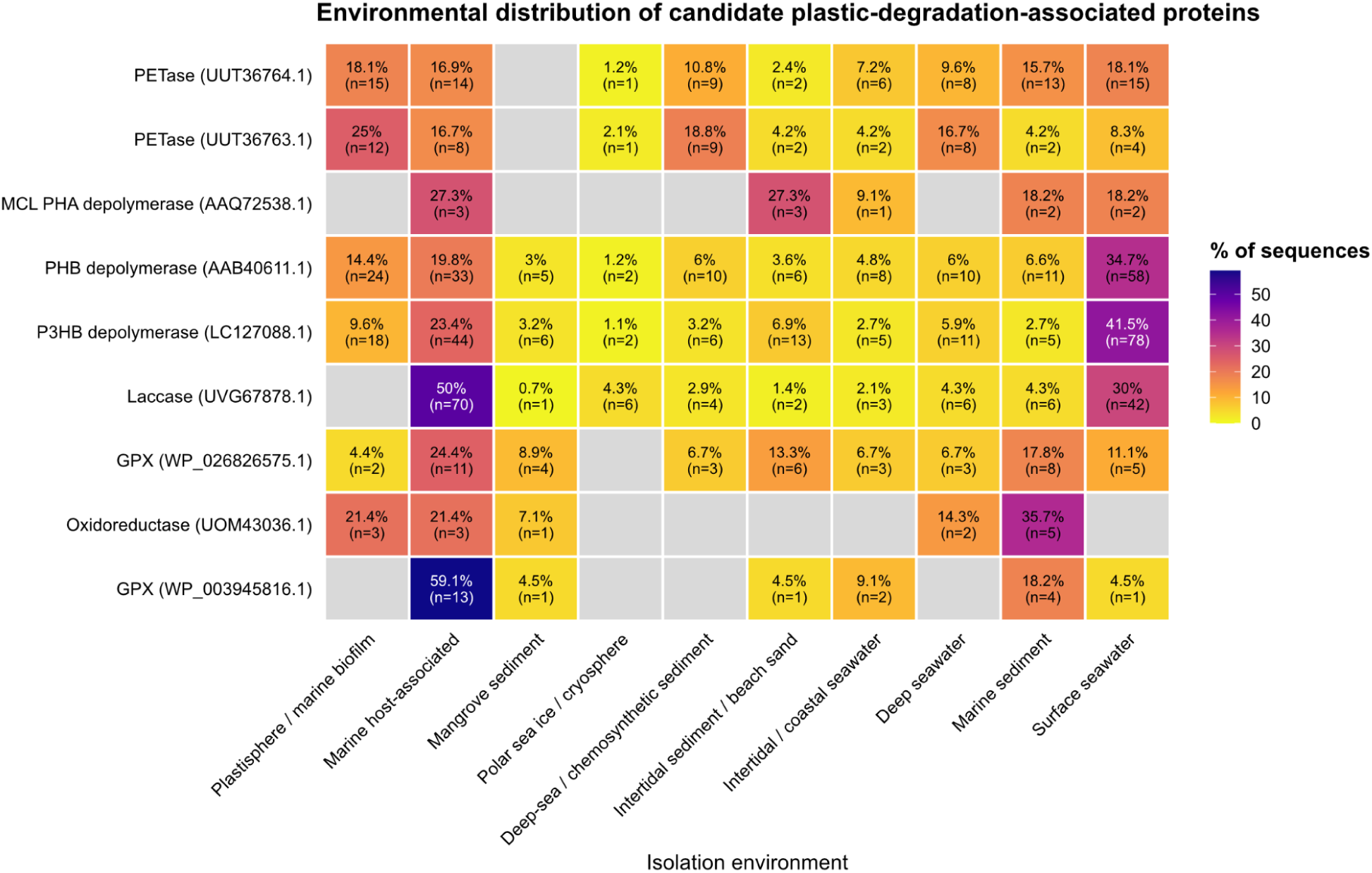
Heatmap showing the environmental distribution of candidate plastic-degradation-associated protein homologs. Rows represent the nine candidate protein groups, and columns represent environmental categories. Values within cells indicate the percentage (top) of sequences within a tree and the corresponding number (bottom) of representatives; gray cells indicate no sequences represented in that environmental category.

Although the homologous proteins are broadly distributed, the strength of the plastisphere signal differs considerably among enzyme families. Across all nine datasets, plastisphere or marine-biofilm sequences appeared in six, with PETase (UUT36763.1) having the largest fraction of representative homologs (12/48, 25.0%). Next was oxidoreductase (3/14, 21.4%), followed by PETase (UUT36764.1; 15/83, 18.1%), PHB depolymerase (24/167, 14.4%), and P3HB depolymerase (18/188, 9.6%). In comparison, plastisphere/marine biofilm sequences represented only 4.4% of the glutathione peroxidase (WP026826575.1) dataset (2/45), and none were seen in the PHA depolymerase, laccase, or glutathione peroxidase (WP003945816.1) datasets.

**Fig. 5** also shows large variations in environmental breadth. Both the PHB (AAB40611.1) and P3HB (LC127088.1) depolymerase datasets included all 10 environmental categories, whilst the PETase (UUT36763.1), PETase (UUT36764.1), laccase (UVG67878.1), and glutathione peroxidase (WP026826575.1) each included nine. Meanwhile, the glutathione peroxidase (WP003945816.1) had six, and the PHA depolymerase (AAQ72538.1) and oxidoreductase (UOM43036.1) datasets each contained five. Several datasets were strongly weighted toward a single environmental category. For example, marine host-associated accounted for 70 of 140 laccase (UVG67878.1) sequences (50.0%) and 13 of the 22 glutathione peroxidase (WP003945816.1) sequences (59.1%), while surface seawater was the largest category for both P3HB depolymerase (LC127088.1; 78/188, 41.5%) and PHB depolymerase (AAB40611.1; 58/167, 34.7%). Within the oxidoreductase (UOM43036.1) dataset, 5 out of 14 (35.7%) accessions belonged to isolates from marine sediment.

## Discussion

This research combines comparative sequence analysis and phylogenetic reconstruction to investigate the taxonomic and geographic diversity of marine microorganisms with candidate plastic-degradation-associated proteins. By extracting marine microorganisms with protein sequence records from PlasticDB (16) and identifying homologous genes from NCBI, we curated a list of putative plastic-degrading microorganisms and their enzymes. We also constructed phylogenetic trees of these taxa, along with models of their reported geographic distributions and isolation environments, to assess whether potential plastic-degrading genes are common within specific evolutionary lineages and marine areas.

### Candidate plastic-associated metabolic potential extends beyond PET hydrolase homologs

Our nine query-derived datasets broaden the study beyond homologs of canonical PET hydrolases and are best interpreted as distinct classes of a candidate plastic-associated metabolic repertoire. The PETase (UUT36764.1 and UUT36763.1)-associated homologs are the strongest candidates for polymer-hydrolysis assays, whereas the MCL PHA (AAQ72538.1), PHB (AAB40611.1), and P3HB (LC127088.1) depolymerase-associated homologs connect the survey to natural polyester metabolism (49, 50, 52). Oxidoreductase (UOM43036.1) and laccase (UVG67878.1) homologs may participate in oxidative transformations, and glutathione peroxidase (WP026826575.1 and WP003945816.1)-associated homologs are more plausibly related to redox protection and persistence in biofilms (53–56). These datasets therefore distinguish proteins that may transform polymers or oligomers from accessory proteins that may support growth on plastic surfaces. These groups encompass hypotheses involving direct polymer hydrolysis and accessory physiological roles, but each requires family-specific experimental validation.

Among the enzyme families examined, analyzing the recovered PETase-associated homologs for synthetic aromatic-polyester cleavage is likely to yield the highest success because characterized PETase provides a structural and catalytic platform for depolymerizing PET and related aromatic polyesters (51). Yet, many enzymes associated with plastic degradation belong to broader protein families with native biological functions unrelated to anthropogenic plastics (52, 57, 58); the MCL PHA and PHB depolymerase families exemplify this, extending to proteins involved in natural polyester metabolism. Polyhydroxyalkanoates are not primarily anthropogenic pollutants but rather naturally occurring classes of microbial carbon- and energy-storage polymers that many bacteria synthesize and later break down, particularly under nutrient-limited conditions (52, 59). Enzymes involved in PHA and PHB turnover may represent polyester-processing scaffolds from which activity may emerge on structurally related synthetic materials. While some previously characterized enzymes act on both natural and synthetic polyesters, this activity cannot be extrapolated to all homologs recovered in this study (59).

In addition to direct ester-bond hydrolysis, the homologs associated with the laccase-like multicopper oxidases and other oxidoreductases could potentially oxidize polymer surfaces and/or weathering products (53, 60, 61). Oxidative modification can introduce oxygen-containing functional groups into recalcitrant polymers or their lower-molecular-weight derivatives, potentially increasing their reactivity and accessibility to subsequent microbial metabolism (24, 62). Laccases have been experimentally associated with polyethylene oxidation and depolymerization (63), while a glutathione peroxidase from a marine Rhodococcus isolate was implicated in low-density polyethylene (LDPE) depolymerization in cooperation with superoxide radicals (24). Glutathione peroxidases may additionally support plastic-associated metabolism by mitigating oxidative stress generated during biofilm growth or on weathered surfaces (54–56). However, these activities are enzyme- and substrate-specific, and sequence homology alone does not establish polymer-degrading activity (57, 58).

Therefore, the nine datasets support a hierarchical approach to experimental testing. PETase- and PHA/PHB-associated hydrolase homologs should be tested directly against defined polymers and oligomers. Oxidoreductase and laccase homologs are better suited to assays of oxidative activity or activity against compounds associated with weathered polymers, whereas the two glutathione peroxidase datasets support tests of oxidative-stress tolerance during plastic surface colonization. This framework keeps accessory physiological traits separate from evidence of direct polymer cleavage while providing candidates for plastic-bioremediation testing.

### Query overlap and family heterogeneity shape phylogenetic interpretation

The nine recovered maximum-likelihood phylogenies harbored significant taxonomic breadth spanning bacterial lineages from narrow clades belonging to phylum Actinomycetota and order Bacillales-associated sequences, to polyester hydrolase-associated phylogenetic clades spanning several Actinomycetota and Gammaproteobacterial genera (**Figs**. **2** to **4**; **Table S1**). The two PETase-associated query searches recovered substantially overlapping α/β-hydrolase sequence space (**Fig. 2**). Recurrence of such PETase-associated homologs suggests that related hydrolytic sequence space is represented within lineages already common in marine surface, sediment, and host-associated settings. This wide range of species mirrors the breadth of the α/β-hydrolase superfamily, which includes many enzyme classes with a conserved structural core (65). In most cases, substrate specificity within this superfamily depends on variable caps, lids, flaps, and accessory domains rather than the conserved core (65).

PHA and PHB depolymerases similarly represent polyester-hydrolyzing enzymes, providing a useful comparison with the PETase-associated datasets. The relatively narrow MCL PHA dataset contrasts with the broader, strongly overlapping PHB- and P3HB-associated datasets, indicating that the three queries partition related polyester-hydrolase sequence space differently. PHA depolymerases vary in sequence and substrate specificity despite commonly sharing an α/β-hydrolase fold and catalytic triad (66). The PHA Depolymerase Engineering Database classified 587 proteins, represented by 735 sequence entries, into eight superfamilies and 38 homologous families (66). This diversity and biochemical classification support analyzing the MCL PHA-associated homologs separately from the two PHB-associated datasets. In comparison, the PHB- and P3HB depolymerase-associated trees contain strongly overlapping sets of Gammaproteobacterial genera (Figs. 3B and C). The overlap between the PHB- and P3HB-associated datasets is expected because PHB and P(3HB) refer to the same short-chain-length PHA (52). The two PHB-related queries may therefore provide redundant samples of candidate plastic-degrading marine homologs. There was no overlap in accessions of oxidoreductase and laccase with both glutathione peroxidase homologs.

These phylogenetic patterns should be interpreted within the constraints of our sequence-based approach. Tree terminals represent protein accessions rather than independent genomes or organisms, and therefore describe sequence relationships within query-defined groups rather than the relative abundance of their microbial hosts. Differences among the PETase-, MCL PHA-, and PHB-associated phylogenies may also reflect the properties of the reference sequences and BLASTP recovery, rather than verified differences in substrate range or evolutionary history. Phylogenetic placement alone cannot establish substrate specificity, independent evolutionary origins of plastic-degrading activity, or confirmation of plastic biodegradation. Instead, these phylogenies provide a framework for identifying phylogenetically diverse and nonredundant candidates for experimental investigation. Biochemical assays will ultimately be required to determine whether individual homologs act on PET, MCL -PHAs, PHB, other polymers, or none of these substrates. Thus, the homologs identified here represent biochemical potential worth investigating rather than evidence of confirmed plastic-degrading activity (57, 58, 64).

### Oxidative- and stress-associated phylogenies support distinct experimental priorities

The laccase (UVG67878.1) homolog phylogeny was among the most taxonomically restricted, with *Psychrobacter* accounting for 128 of 140 source-organism records (**Fig. S4**; **Table S1**). This concentration indicates that the dataset primarily captures multicopper oxidase diversity within a single genus. Structurally, laccases are one functional subtype of the broad superfamily of multicopper oxidases (53), whose members can differ in substrate specificity. Thus, the identified homologs provide a starting point for investigating the functional diversity of these enzymes in *Psychrobacter*, including their potential roles in plastic-associated oxidative chemistry. The observed taxonomic concentration may be due to the query sequence, RefSeq composition, the marine-provenance filter, or their combined effects. Accordingly, gene-copy analysis and gene-tree/species-tree reconciliation would be required to test expansion within *Psychrobacter*.

Studies of polyethylene-associated microorganisms provide useful assays for evaluating laccase-like homologs, while also illustrating the limits of functional inference. *Pseudomonas citronellolis* str. E5 and *Rhodococcus erythropolis* str. D4 exhibited extracellular laccase-like activity during growth with low-density polyethylene powder; however, activity was measured using 2,6-dimethoxyphenol rather than polyethylene itself (60). The highest measured activities occurred with added copper, supporting multicopper oxidase activity without demonstrating polyethylene cleavage. A separate multi-omics study detected a PlasticDB-annotated laccase in an early polyethylene biofilm, but protein detection did not identify the substrate or reaction catalyzed in that community (61). Thus, assays of the present homologs must distinguish oxidation of soluble substrates from chemical transformation of an intact polymer.

The two glutathione peroxidase-associated queries share broad functional annotations but appear to retrieve distinct sequence and taxonomic groups. Many bacterial proteins annotated as glutathione peroxidases are thioredoxin-dependent peroxiredoxins rather than canonical glutathione-dependent enzymes, and reductant specificity depends on sequence and structural features that broad annotations may not capture (54). Deletion of a bacterial glutathione peroxidase increased susceptibility to hydrogen peroxide, hypochlorous acid, and an organic peroxide (55), and oxidative-stress responses can alter biofilm formation and extracellular polymeric-substance (EPS) production under oxidant exposure (56). These findings support comparative tests of peroxide tolerance and expression during matched plastic and inert surfaces. Polymer-cleavage assays would be warranted if structural or biochemical evidence suggests a direct role in polymer transformation.

### Marine plastic-degradation-associated homologs span diverse habitats beyond plastic-associated biofilms

The marine provenance of the results argues against interpreting the recovered homologs as plastisphere-specific. Only 74 of 718 query-to-hit assignments (10.3%) were linked to plastisphere or marine-biofilm samples, while 644 (89.7%) came from other marine categories (**Fig. 5**). This distribution appears more consistent with protein families that participate in wider marine physiological processes than with functions restricted to plastic-associated communities. Differences among datasets are more useful for choosing comparisons than for ranking plastic association. PETase (UUT36763.1) had the largest within-dataset plastisphere fraction at 12 of 48 assignments (25.0%). In contrast, no plastisphere or marine-biofilm assignments were found in the MCL PHA (AAQ72538.1), laccase (UVG67878.1), or glutathione peroxidase (WP003945816.1)-associated datasets (**Fig. 5**). Provenance breadth also supports different biological expectations across protein families. PETase (**Fig. 2**) and PHA-associated homologs (**Fig. 3**) were found in organisms isolated from nearly all environmental categories (**Fig. 5**). Their broad distribution is compatible with established roles in natural-polyester metabolism predating anthropogenic plastic exposure; hence, activity against synthetic substrates remains possible (52, 59). By contrast, marine host-associated samples accounted for nearly half of all laccase (UVG67878.1) and glutathione peroxidase (WP003945816.1)-associated assignments. This concentration of marine-host-associated samples motivates tests of multicopper-oxidase and redox-protection functions in host-associated and surface-attached growth (53–55).

Evidence from controlled plastisphere studies further reinforces this distinction between provenance and enrichment. A meta-analysis of 2,229 plastisphere samples from 35 studies found that environmental and study-design variables strongly influenced community composition and could outweigh the effect of plastic type (67). In a paired field experiment, biofilms on PE, PA/nylon, and glass differed from surrounding seawater but not significantly from one another overall, suggesting a substantial contribution from general surface colonization (68). A recent synthesis likewise argued that claims of plastisphere uniqueness and plastic biodegradation require comparative designs and direct evidence beyond colonization (69). Matched plastic and nonplastic surfaces could test enrichment; polymer-specific product formation could test polymer transformations; and carbon assimilation, potentially through the use of DNA- or RNA-stable carbon isotope probing, could address utilization (69–73).

Despite the geographic distribution map representing a nonstandardized biogeographic sample, our spatial visualization expands the range of candidate comparisons. Mappable locations were available for 475 of the 718 query-to-hit assignments, representing 337 distinct accessions, and included coastal waters, deep seawater, sediments, host-associated samples, mangroves, and polar settings (**Fig. 1**; **Table S1**). Deep-water and sediment accessions are relevant to future sampling because turbidity currents can transport and bury microplastics in deep-marine sediments, and the deep sea is a documented sink for microplastic debris (74, 75). Our dataset compiles published isolates and their NCBI records; consequently, research effort becomes part of the signal we measure. The number of times a region contributes candidate sequences to public databases reflects the extent of marine microbial plastic research conducted in that region, regardless of whether it truly harbors a disproportionately high number of plastic-degrading enzymes. As a result, the apparent biogeographic hotspots and the wide range of habitats of candidate homologs reinforce the possibility that these genes perform other natural biochemical functions while their plastic-related activities remain untested.

## Conclusions

Marine microorganisms represent a largely untapped source of proteins with potential roles in plastic transformation, offering opportunities to discover enzymes and biochemical processes beyond those already characterized. To explore this potential, we combined PlasticDB and NCBI BLASTP homology searches to identify candidate plastic-degradation-associated proteins from marine microorganisms. Using nine curated reference protein sequences of marine origin, representing PETase-, laccase-, MCL PHA/PHB depolymerase-, oxidoreductase-, and glutathione peroxidase-associated families, we identified 718 query-to-hit assignments (523 unique protein accessions). We then used phylogenetic reconstruction and geographic mapping to investigate the evolutionary relationships and environmental distributions of these candidate homologs.

Overall, our curated provenance metadata and phylogenies produce an experimentally tractable set of marine protein homologs. Our study sorted a large genetic dataset into four testable biological roles: synthetic-polyester hydrolysis, natural-polyester metabolism, oxidative chemistry, and redox protection during surface-associated growth. With query overlaps reducing the number of independent candidate pools, future lab testing can be more efficient by pinpointing phylogenetically separated lineages that retain meaningful sequence differences.

However, several cautions are warranted to avoid overinterpreting claims that these patterns reflect a true biological signal. At the sequence level, homology to a known plastic-degrading enzyme may indicate structural relatedness, but does not establish that the candidate in question is itself a catalyst capable of degrading plastics (57, 58); enzyme families such as the PHA/PHB depolymerases have well-documented native roles in microbial carbon storage, which are unrelated to synthetic plastics (52, 66). Likewise, the apparent biogeographic hotspot may reflect uneven regional sampling and coordinated reporting rather than a true ecological hotspot of plastic degradation, and the wide range of habitats of candidate homologs reinforces the possibility that these genes perform other natural biochemical functions while their plastic-related activities remain untested.

Future work should move from sequence potential to mechanism and ecological relevance in stages. Family-appropriate biochemical assays should first compare plastisphere-derived proteins with close nonplastisphere homologs from separated branches and quantify defined polymer-derived products. Genomic-context analyses (74), together with structural features, secretion predictions, copy number (71), and gene-tree/species-tree reconciliation, can then refine mechanistic and evolutionary hypotheses (76). Finally, matched sampling of plastic, inert surfaces, water, and sediment, combined with metagenomic, transcriptomic or proteomic, product-specific chemical, and isotope-tracing measurements, would connect candidate presence to enrichment, expression, polymer transformation, and carbon assimilation. This sequence of tests offers a direct route from the present candidate shortlist to experimentally supported marine plastic-biotransformation pathways.

The findings together produce a prioritized, taxonomically and geographically diverse shortlist of putative plastic-degrading enzyme homologs suitable for future experimental validation, alongside methodological guidance for future research, including targeting underrepresented ocean basins and prioritizing lineages already associated with these gene families. As biotechnological approaches to plastic bioremediation continue advancing, comparative studies such as this one may help direct limited experimental resources toward the microbial taxa and marine regions most likely to yield validated, plastic-degrading enzymes.

## Data availability

NCBI RefSeq and PlasticDB are openly accessible resources. Accession-level sequence and provenance metadata are provided in **Table S1**. FASTA files, sequence alignments, complete and displayed phylogenetic trees, environmental-distribution outputs, and analysis scripts are available at https://github.com/corporeal-snow-albatross-5/Marine_Microbial_Plastic_Degradation_Potential. Geographic mapping code and standardized geographic datasets are available at https://github.com/chloekong0110/Candidate-Plastic-active-Enzyme-Homologs-Map.

## Funding information

This research received no specific grant from any funding agency in the public, commercial, or not-for-profit sectors.

## Conflict of interest

The authors declare no conflict of interest.

## Author contributions

CK: Investigation, Data curation, Methodology, Validation, Visualization, Formal analysis, Writing - original draft, Writing - review & editing. SE: Conceptualization, Investigation, Supervision, Data curation, Methodology, Validation, Visualization, Formal analysis, Writing - review & editing.

